# Assessing computational strategies for the evaluation of antibody binding affinities

**DOI:** 10.1101/2025.09.25.678535

**Authors:** Ida Autiero, Damiano Buratto, Fengyi Guo, Wanding Wang, Malay Ranjan Biswal, Kevin C. Chan, Ruhong Zhou, Francesco Zonta

**Affiliations:** Department of Biomedical Sciences, Institute of Biostructures and Bioimaging, National Research Council (CNR), Napoli, Italy; Institute of Quantitative Biology, Shanghai Institute for Advanced Study, College of Life Sciences, Zhejiang University, Hangzhou, China; Department of Biosciences and Bioinformatics, School of Science, Xi’an Jiaotong-Liverpool University, Suzhou, 215123, China

## Abstract

Accurate evaluation of binding affinity is critical in drug discovery to identify molecules that bind strongly to their targets while minimizing off-target effects. Although binding affinity calculations are theoretically well-defined, they require exhaustive sampling of configurational space, a step that often requires significant computational resources. In this study, we compare different methods for calculating the binding energy of antibodies targeting a peptide derived from the N-terminus of CXCR2, a GPCR-family protein. Contrary to some previous reports, we find that equilibrium MMPBSA calculations yield better agreement with experimental binding affinities than non-equilibrium potential of mean force evaluations, underscoring the system-dependent performance of these methods. We also observed a modest improvement in accuracy when MMPBSA is combined with replica exchange molecular dynamics, albeit at a significantly higher computational cost. Calculation based on Rosetta force field, instead, produced results that did not correlate with the experimental data. We attribute these findings to two factors, which could limit the applicability of some methodologies that are widely used in the computation the binding energy: the high potency of the antibodies studied and the dominance of hydrophobic interactions between the antibodies and the peptide. Overall, this work provides important insights for optimizing *in silico* antibody screening strategies.

## 1. Introduction

The relevance of computer-aided drug design has increased in recent years, together with the availability of computational power and the rapid growth of machine learning methods applied to computational biology [1–3]. Computational prediction of experimental results can diminish the time and the cost of developing new drugs and may revolutionize the field of drug discovery [4]. Among the possible applications of computational methods, the calculation of binding affinity is of primary interest, as drugs with high affinity to a specific target can be used in lower concentration, minimizing the risk of unwanted interactions with other targets [5]. Similar calculations can also be used in understanding pathologies, for example, to evaluate how mutations can affect protein-protein interactions [6].

The recent development of highly successful computational methods for de novo protein binder design, such as BindCraft [7] and Rosetta Fold diffusion [8], which can generate hundreds of candidate binders to a chosen target, represents a major breakthrough but does not eliminate the need for rapid and accurate evaluation of binding affinities through computational methods. Generative methods, indeed, cannot rank their prediction in an efficient way, and false positives are common.

Machine learning methods for rapid evaluation of binding affinities have also emerged in recent years, but they mostly rely on training on experimental or predicted datasets [9–11]. Therefore, they may lack generalizability to out-of-distribution examples [12] and are not always transferrable between different molecular systems.

Another limitation of machine learning methods is that they often rely on a single configuration to predict the binding of the two molecules. In reality, the interaction between two proteins (or a protein and a small molecule) is very dynamic, and the energies should be, at least in principle, evaluated on many different configurations sampled from the correct ensemble.

Configuration sampling and binding affinity calculations can be obtained from molecular simulations [13,14]. However, such calculations are often computationally prohibitive as they require the sampling of the very vast configuration space the molecular system lives in, and this issue is exacerbated in the case of protein-protein interactions.

For this reason, many different simplified algorithms have been proposed (for example [15–19]), with various degrees of approximation. However, no “golden rule” has emerged on the best strategy for calculation of binding affinities between two proteins, likely because the performance of these methods could be system dependent [20].

In this study, we explore the efficacy of various strategies based on molecular dynamics (MD) to evaluate the binding affinity of an antibody to its target and will compare this with previously published results. In this context, we want to obtain (i) a sensitive sampling of the configuration space, and (ii) an accurate calculation of the binding energy for each configuration.

The first approach we consider here is the MMPBSA (Molecular Mechanics Poisson Boltzmann Surface Area) method [15], which estimates the Gibbs free energy as a sum of internal and solvation energy components. The binding free energy is then obtained as the difference between the Gibbs free energies in the bound and unbound state. We have previously shown [20] that this method produces a reliable prediction on our chosen dataset (see below). In this work we want to test whether the results can be improved with an enhanced sampling of the conformational space. To this purpose, we paired MMPBSA calculation with replica exchange molecular dynamics at different temperatures (T-REMD) [21], while imposing positional restraint to the proteins outside the interaction interface, to keep the computational costs manageable.

We will then compare the binding energy determination obtained with the MMPBSA method with that obtained by the Rosetta energy function [22] on the same dataset.

Finally, we will consider a completely different approach which evaluates the binding free energy from the out of equilibrium Potential of Mean Force (PMF), i.e. the average work required to physically separate two interacting molecules [23]. Typically, the PMF is computed along a reaction coordinate which corresponds to the spatial separation between the two molecules with some sampling strategies such as umbrella sampling [24,25].

The system chosen is the complex formed by a potent anti CXCR2 antibody and a peptide derived from the N-terminus of the CXCR2 [26] (**Figure 1 panel A**). The antibody displays a peculiar binding with its peptide epitope, as the interaction is stabilized by a hydrophobic pocket found between the heavy chain and the light chain complementary determining regions (CDRs) [20]. In addition to the wild-type antibody, we evaluate the binding of eight of its variants to the same target, covering approximately a tenfold range in experimentally measured binding affinity.

**Figure 1.**
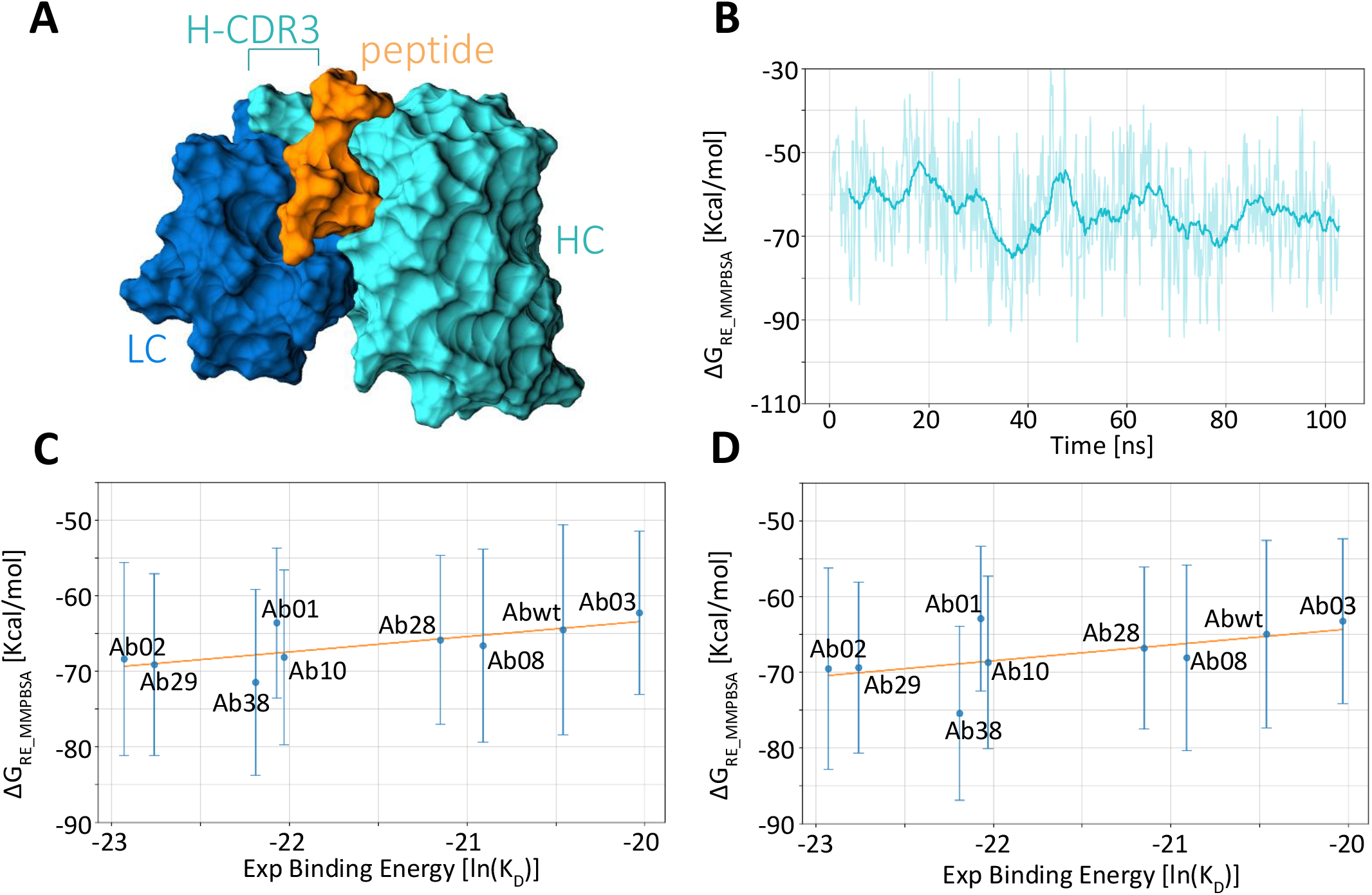
Calculation of different antibodies binding energies to the peptide. (A) Representation of the antibody – peptide complex. Only the variable part of the antibody (VH and VL region) is taken into consideration during the simulation. The antibody heavy chain is shown in cyan and the light chain in dark blue. The peptide is represented in orange color. (B) Values of Δ*G*for the complex formed by the Abwt and the peptide for all the frames of the 300 K replica. Values are shown as raw data (thin line) and their running average (thick line). Panels (C) and (D) show the correlation between the calculated binding free energies and their respective experimental values. The results were obtained using the whole 20-100 ns trajectories (C) or the shorter 20-50 ns trajectories. The correlation coefficients are R^2^ = 0.31 and R^2^ = 0.57, respectively.

## 2. Results

### 2.1 Effects of sampling on binding free energy calculation

In our previous works [20,27], we aimed at obtaining a reliable estimation of the binding free energy in a relatively short computational time. Here, we investigate whether we can enhance the accuracy of our prediction by applying alternative strategies to obtain a more accurate sampling of the configuration space.

The first strategy explored is the use of T-REMD [21]. In this approach, multiple simulations (replicas) are run at different temperatures. The replicas can exchange temperature at regular intervals, allowing lower temperature simulations to explore a larger configuration space.

For each of the nine antibody variants considered, we performed 64 replicas of 100 ns simulations, with temperatures ranging from 300 to 380 K, resulting in a total simulated time of 57.6 μs. We then used the MMPBSA method to calculate the binding affinity of each antibody to the peptide antigen using frames extracted from the replica at 300 K (**Figure 1 panel B**). Sampling enhancement was verified by monitoring the Ramachandran angles for the peptide residue critical to binding (**Supplementary Figure 1**): the exploration of the φ–ψ torsion angles is noticeably greater in the T-REMD simulations than in the standard MD simulations.

Analysis of the variation of binding free energy vs time for the ensemble of all the replicas shows that, in most cases, the free energy decreases during the first 20 ns of the simulation before reaching a stable plateau (**Supplementary Figure 2**). This appears to be the time required to obtain an appropriate number of exchanges between the different temperature replicas i.e. to achieve equilibration for the 300K simulations. Therefore, the first 20 ns were excluded in the calculation of the average binding free energies.

Moreover, the calculated binding free energy shows little dependence on temperature, as simulations at different temperatures yield similar results. This is consistent with the binding mode observed in our simulations, where the peptide is deeply buried in a hydrophobic pocket of the antibody. This confinement restricts the peptide’s conformational space, leading to a small conformational entropy change upon binding and thus a weak temperature dependence.

In our previous work, longer simulations did not necessarily yield better agreement with experimental data. This trend was confirmed also here, where the prediction based on the 20-50 ns interval showed a higher correlation (R^2^ = 0.57) with the experiments than the prediction obtained with the full 20-100 ns interval (R^2^ = 0.29) (**Figure 1 panels C, D**).

The result based on the 20-50 ns interval, correlates as well with experiments as our previous estimate (R^2^ = 0.57), but it must be noted that previous results were obtained with a much-reduced sampling [20]. However, the binding affinity difference calculated using the replica exchange is more realistic than in our previous estimation, having obtained a maximum ΔΔ*G*around 6 Kcal/mol instead of unrealistic differences of more than 10 Kcal/mol obtained in our earlier work. **Figure 2** shows the frame-by-frame MMPBSA estimates of binding free energy from the 300K replica for each antibody variant confirming that after the first 20 ns, simulations are generally stabilized.

**Figure 2.**
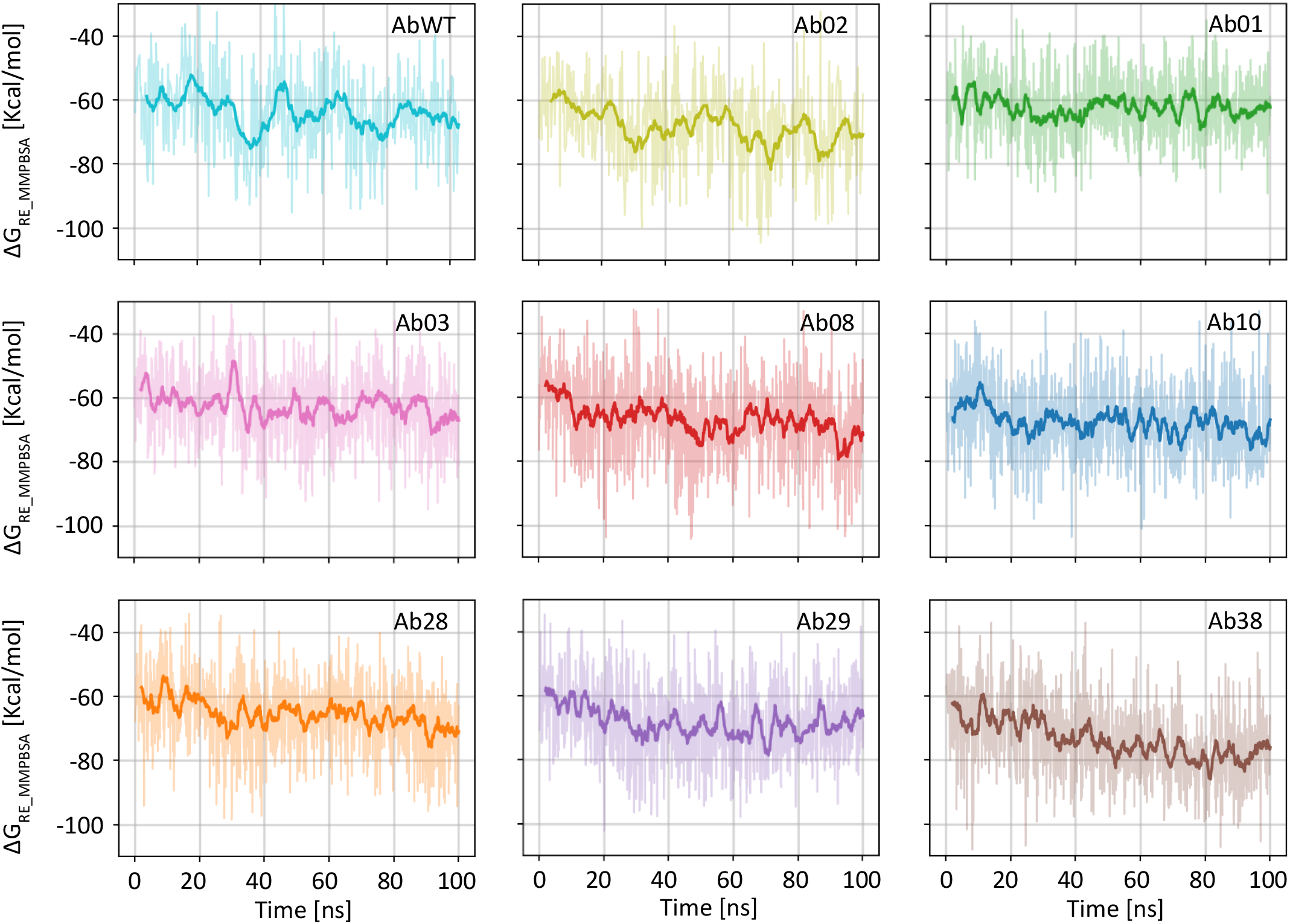
Full dataset for the binding energy calculation of all antibody variants using the MMPBSA method. Each graph shows the computed binding free energy, calculated on the 300K replica for one of the antibody variants. The data for AbWT are the same as Panel B of Figure 1. The energy trend is different in various cases, with some systems equilibrating faster. However, the first 20 ns tend to show a higher average than the rest of the simulation and therefore have been discarded in the following analysis. Values are shown as raw data (thin line) and their running average (thick line).

Finally, we can observe that MMPBSA calculation provides better estimates than predictions made with Rosetta using the same configuration dataset. The correlation with experiments is indeed R^2^= 0.01 for the 20-50 ns set and R^2^ = 0.02 for the 20-100 ns when we use the Rosetta energy function to calculate the binding energy (**Figure 3**). Furthermore, contrary to the calculation performed with the MMPBSA method, where transient trends may appear on timescales of about 10 ns, binding energies calculated with the Rosetta energy function never change significantly during the simulation, indicating that this calculation method is less sensitive to minor configurational variations.

**Figure 3.**
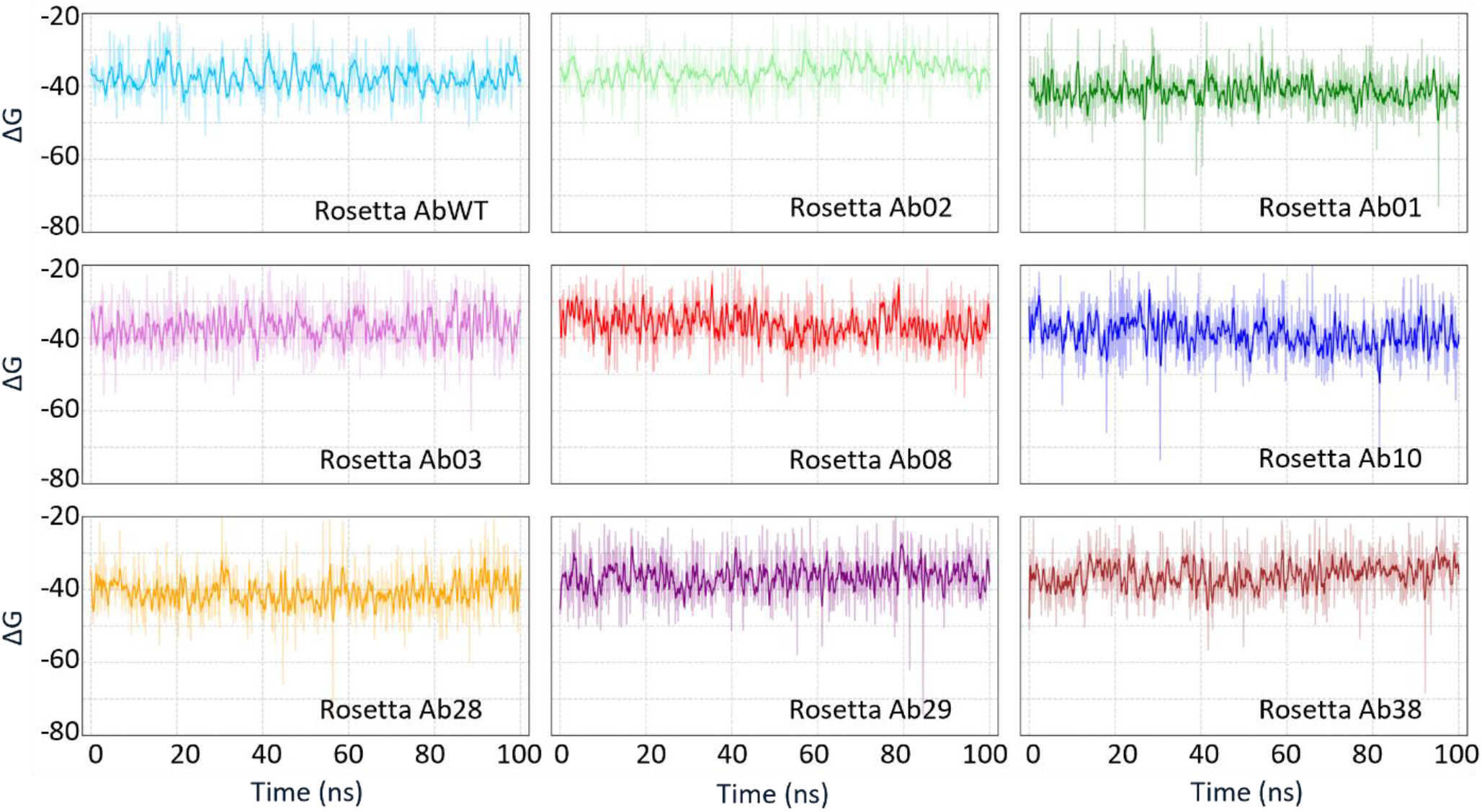
Full dataset for the binding energy calculation of all antibody variants using Rosetta energy function. Each graph shows the binding energy computed using Rosetta for a different antibody bound to the peptide. Configurations are taken from the 300 K replica, as in figure B. Energies calculated with Rosetta show smaller fluctuations and there are no appreciable trends on longer timescales. Values are shown as raw data (thin line) and their running average (thick line).

### 2.2 Molecular mechanisms leading to binding energy differences

Figure 4 shows the representative structures of the complexes of two different antibodies (AbWT and Ab02 – see methods for antibodies naming) and the peptide extracted from the 20-50 ns of T-REMD trajectories. Although the binding modes are globally comparable among these two systems, moderate variations can be appreciated. Replacing serines with arginines in the CDR3 of the antibody, indeed, results in a different orientation of the TYR108 and TYR110 side chains. Both these residues direct the phenolic ring that carries the oxhydryl group towards the solvent, in a parallel and coordinated fashion with ARG106, which exposes its positively charged group to the water. In the WT system, this ordered displacement does not exist. Moreover, the peptide N-terminal region is slightly further away from the antibody surface in the WT system compared to Ab02.

Overall, although it is difficult to rationalize these differences and understand how they can lead to a binding affinity difference, simulations demonstrate their ability to infer important structural insights at atomistic level of recognition and correctly discriminate between good binders and bad binders.

**Figure 4.**
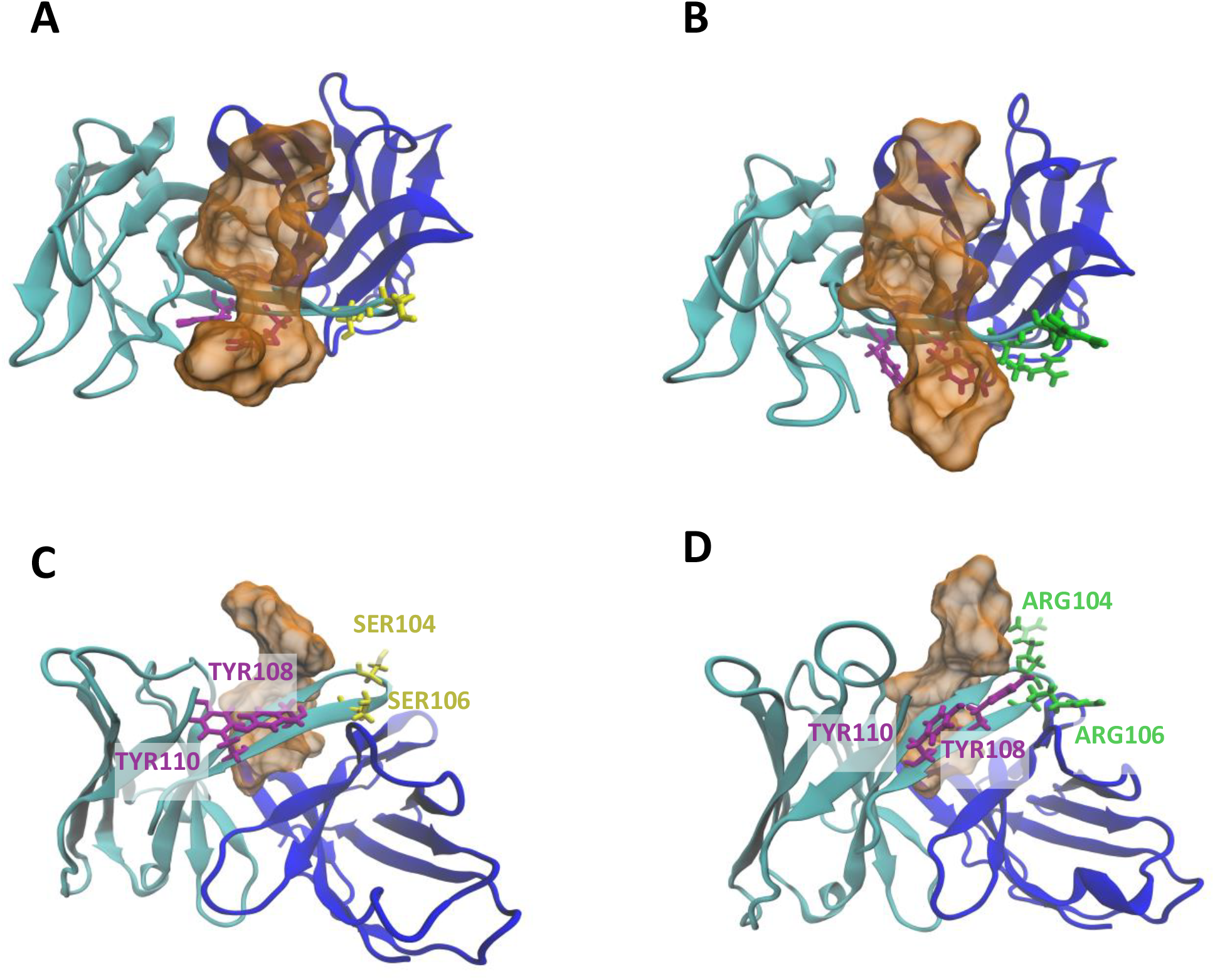
Details of molecular interaction between the antibody and the peptide. (A) and (B): Top view of the REMD representative structures of wildtype and Ab02, respectively. (C) and (D) side view of the same configurations. Representative structures are taken from the average configuration along the trajectories. In Ab02 two serines in CDR3 are replaced by two arginines. This causes a change in the coordination of TYR108 and TYR110 in the same CDR. The color code is consistent with Figure 1.

### 2.3 Evaluation of the potential of mean force with the umbrella sampling method

The PMF is the average of the out of equilibrium work (*W*) necessary to separate two interacting molecules. A common strategy to compute the PMF is the umbrella sampling method, in which the two molecules are kept at a fixed distance by an external force so that their interaction can be measured as a function of the distance along a preferred reaction coordinate [24]. In this approach, artificial biasing potentials are applied to keep the system at fixed values of the reaction coordinate. This forces the molecules into configurations that would be rarely sampled under thermodynamic equilibrium, allowing the exploration of otherwise inaccessible regions of phase space. Each biased simulation produces an estimate of the free energy cost of maintaining the system in that configuration. By combining and reweighting these biased distributions, the effect of the artificial potentials is removed and the unbiased PMF is reconstructed, from which the binding free energy can be extracted as the difference between the bound and unbound states.

In the first iteration of the PMF calculation, we used a fixed 0.5 Å interval between every umbrella sampling windows along the reaction coordinate (the distance between the antibody and the peptide, see methods). However, this spacing proved to be insufficient in most cases, leading to abrupt transitions between the bound and the unbound states and insufficient sampling of intermediate configurations.

Eventually, estimation of the PMF for each antibody was done by setting the window width to 0.3 Å, resulting in approximately 100 umbrella windows per system. For each window, a 2 ns simulation was carried on for a total of about 200 ns for each antibody (∼1.8 μs for the whole set of simulations, **Figures 5 and 6**). The correlation between experiments and simulations in this case is R^2^ = 0.19, notably lower than the results obtained with the MMPBSA method.

**Figure 5.**
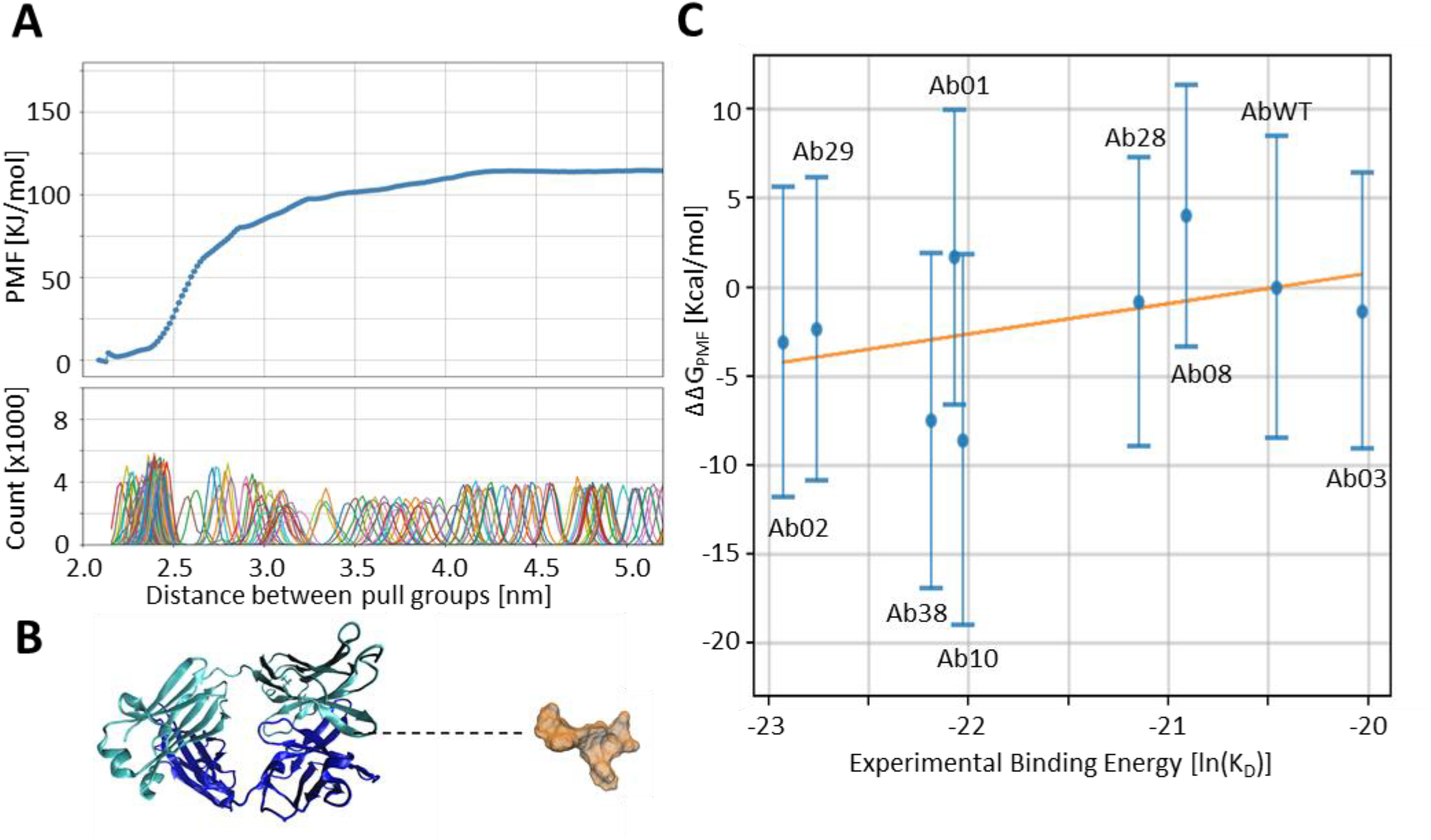
Calculation of the PMF for evaluating the binding energy. Panel A shows the reconstructed PMF profile with the error bars for each point along the reaction coordinate. The PMF has been generated from umbrella sampling along the reaction coordinate. The bottom panel shows the gaussian distributions of the sampled configurations in each window. Panel B is a snapshot of the Steered MD simulations showing the peptide pulled far away from the antibody. The reaction coordinate, i.e. the distance between the two pull groups, is shown as a dotted line, and corresponds to the x-axis of Panel A. Panel C shows the correlation between the calculated binding energy for the different variants using the umbrella sampling method and their respective experimental values.

**Figure 6.**
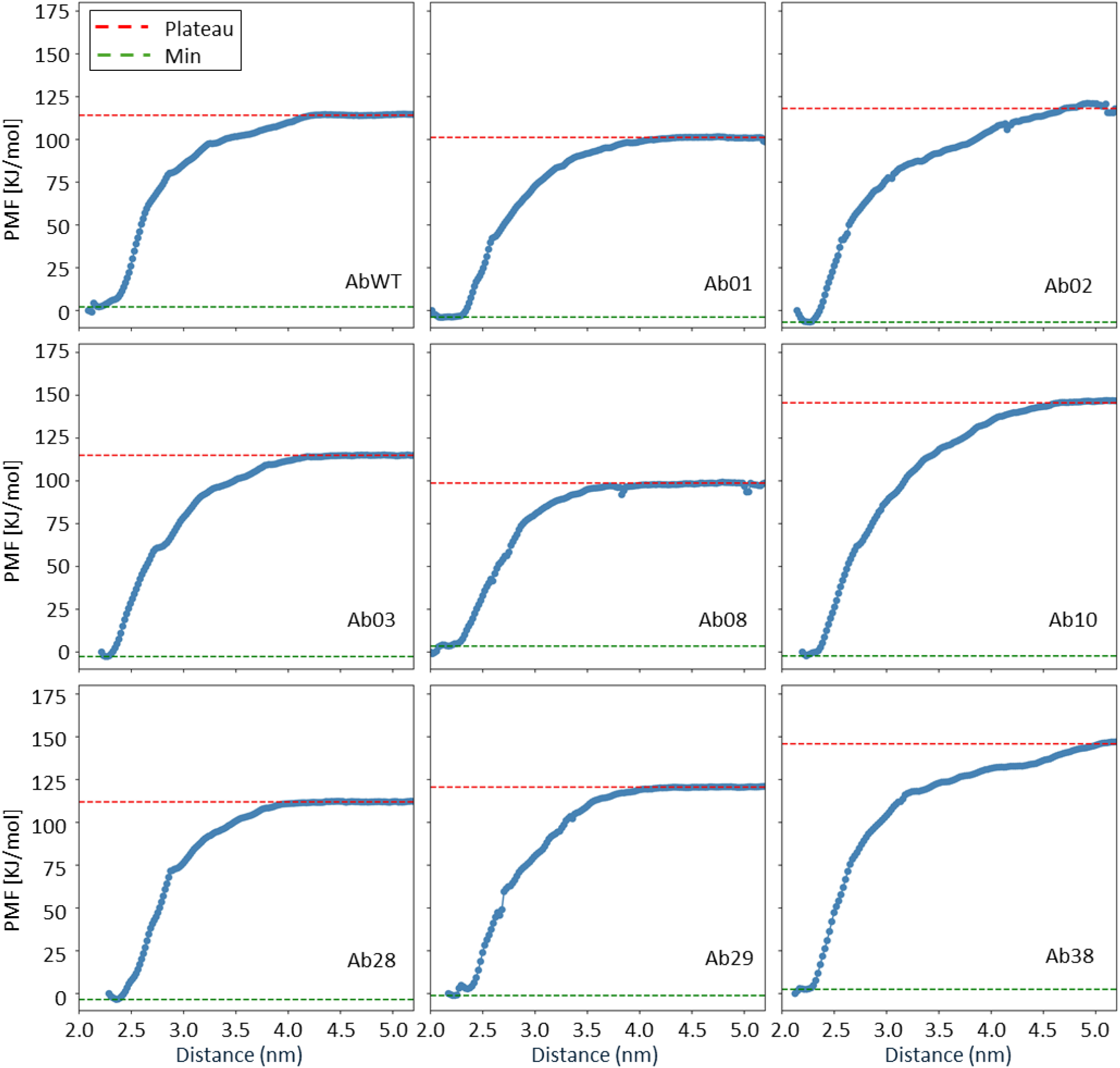
Full dataset for the binding energy calculation of all antibody variants using PMF. Each panel represents the PMF, calculated along the reaction coordinate, for separating a different antibody from the peptide. Higher plateau values correspond to stronger binders, as the work done to separate the two molecules is higher.

### 2.4 Comparison between the computational costs and efficacy of different methodologies

The Replica Exchange MMPBSA (RE-MMPBSA) method has shown the highest correlation with experimental data when considering the 20-50 ns simulation interval. On the other hand, the correlation between different methods is generally low, the best agreement being observed between RE-MMPBSA and umbrella sampling (R^2^ = 0.47, **Table 1**).

**Table 1.** Computed ΔΔG values for each dataset (Kcal/mol). For the experimental determination we used the formula 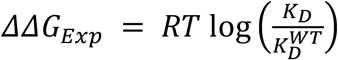 where T is the temperature (300 K) and R is the gas constant.

| | $\Delta\Delta G$<br>RE-MMPBSA<br>(20-50) | $\Delta\Delta G$<br>RE-MMPBSA<br>(20-100) | $\Delta\Delta G$<br>MMPBSA | $\Delta\Delta G$<br>PMF | $\Delta\Delta G_{Exp}$ | $\Delta\Delta G$<br>RE-Rosetta<br>(20-50) | $\Delta\Delta G$<br>RE-Rosetta<br>(20-100) |
| --- | --- | --- | --- | --- | --- | --- | --- |
| Ab01 | 0.9 | 1.2 | -18.4 | 7.0 | -0.96 | -3.50 | -3.32 |
| Ab02 | -4.9 | -4.6 | -29.4 | -12.9 | -1.47 | 0.59 | 2.10 |
| Ab03 | 2.4 | 0.4 | 1.4 | -5.6 | 0.26 | 0.96 | 1.03 |
| Ab08 | -2.4 | -3.9 | -28.7 | 16.8 | -0.27 | 2.12 | 1.21 |
| Ab10 | -4.5 | -4.7 | -33.7 | -35.9 | -0.94 | -0.08 | -1.15 |
| Ab28 | -1.3 | -2.2 | -22.5 | -3.4 | -0.41 | -3.71 | -3.05 |
| Ab29 | -4.6 | -4.7 | -41.3 | -9.8 | -1.37 | 0.33 | 1.36 |
| Ab38 | -6.3 | -10.2 | -15.5 | -31.4 | -1.03 | 0.60 | 1.90 |

Despite these important differences, we can observe how the different methods generally agree in discriminating between strong and very strong binding antibodies, with remarkable sensitivity, even if the experimental difference is relatively small (about 10-fold). For instance, the binding free energy Δ*G*of Ab02 is consistently lower than the Δ*G*of WT, in agreement with experimental observations. Only calculations based on the Rosetta energy function are not sensitive enough in the binding affinity range explored in our systems.

As the maximum span of predicted ΔΔ*G*values presents large differences between different computational methods, to better visualize this result, we consider the normalized value of the binding energy ΔΔ*G* by dividing it by the absolute value of the ΔΔ*G* between the WT and the Ab02, which is considered the best antibody from the experimental point of view:

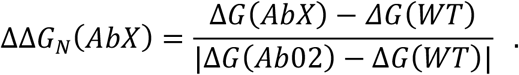

With this definition, the normalized binding energy of the wild-type antibody (ΔΔ*G*_*N*_(*WT*)) and of the Ab02 antibody (ΔΔ*G*_*N*_(*Ab*02)) are zero and minus one, respectively, in each set of data, independently of the computation method. Correlations of the different ΔΔ*G*_*N*_ sets across different computational methods are not affected by this normalization. Experimental values were also normalized using the same approach, after converting dissociation constants *K*_*D*_ into binding free energies using the relation ΔΔ*G*_*N*_(*AbX*) = *RT* In *K*_*D*_(*AbX*), where T is the temperature (300 K) and R is the gas constant.

Figure 7 shows the normalized binding energies computed by the various methods plotted against the same quantity calculated on experimental values. We can notice some regular patterns on the predicted properties of the antibodies, for example, all methods agree that besides the already mentioned Ab02, also Ab29, Ab38, Ab10, Ab28 have a better binding affinity to the peptide than the WT antibody, in agreement with the experiments. This shows that MD-based methods are reliable tools for affinity screening of antibodies.

However, in disagreement with experiments, we observe that simulations almost always identify Ab01 as a worse antibody than the WT. With the same criterion, we can point out two antibodies (Ab10 and Ab38) that may be better than Ab02.

**Figure 7.**
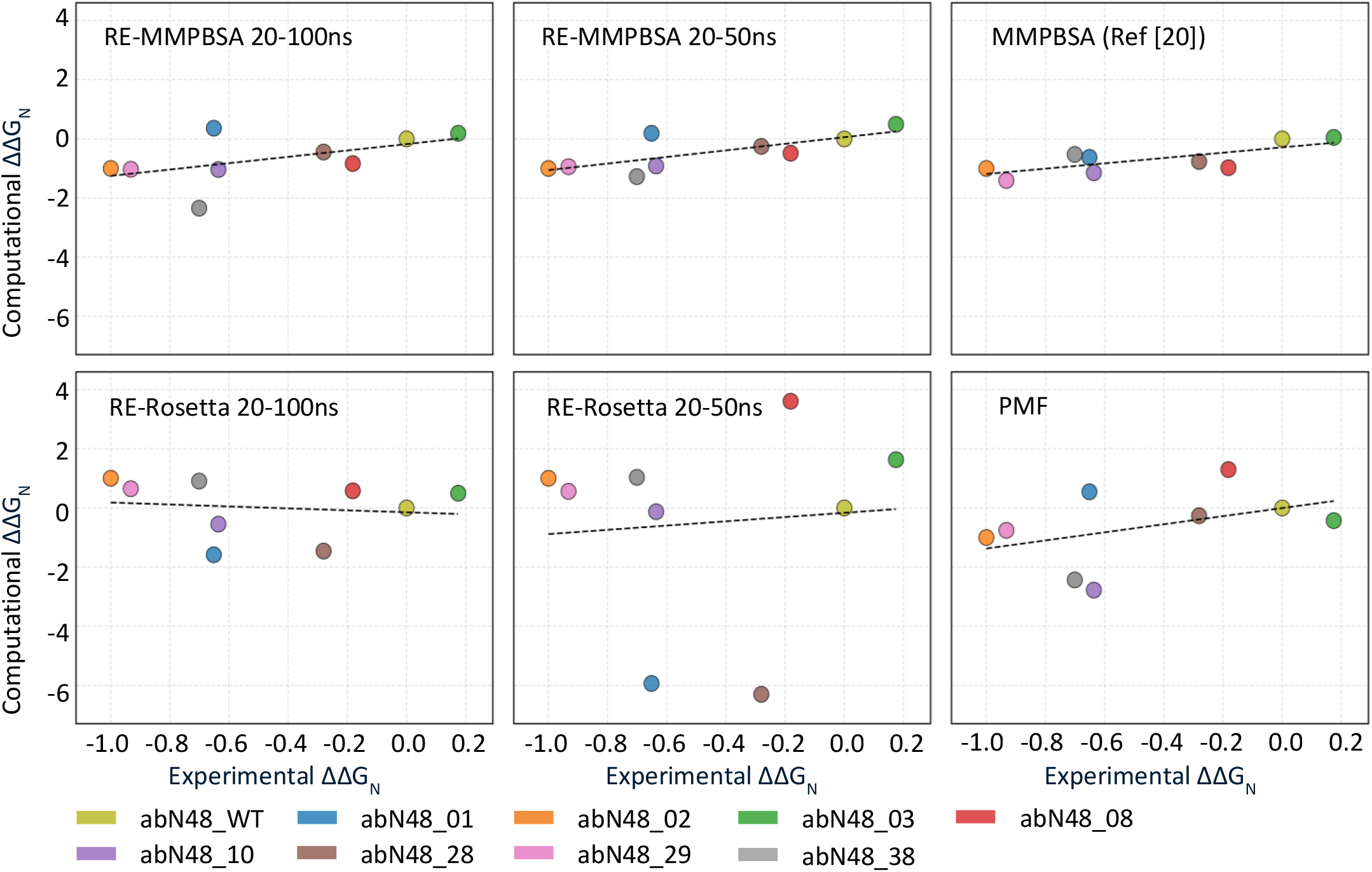
Normalized computational ΔΔ*G* of the various methods vs the experimental values. Each graph reports the ΔΔ*G*_*N*_ obtained with one computational method against the experiments. The best performing methods are based on equilibrium sampling on the configuration space and the use of MMPBSA for energy calculation.

This can be interpreted in two different ways. On one hand, it could indicate a limitation of the computational methods, which failed to correctly sample the configuration space for these specific antibodies. On the other hand, experimental data are also affected by measurement errors, meaning these antibodies may not have been properly characterized in the experiments.

## 3. Discussion

Computation of binding free energies between molecules has important impacts in understanding biological processes and in drug design. However, it remains a difficult task in many practical cases as the existing computation methods cannot always produce reliable estimates. Despite the development of simplified energy functions in the past [18,22,28], and more recent advances in machine learning, which show promise in integrating structural and sequence data [29,30], accurate sampling of the configuration space remains indispensable for binding affinity calculations. This is due to the significant contributions of entropy, the intrinsic flexibility of proteins, and solvent effects. Recent hybrid approaches combining machine learning and molecular simulations [31,32] offer potential pathways to address these challenges but require further development.

In the work presented in this article, we have explored the ability of different methodologies and sampling strategies to reproduce the experimental results of the binding affinity of a series of antibodies to a peptide taken from the N-terminal region of the CXCR2 protein.

In particular, we compared: (i) simple MMPBSA calculations (published in [20]), (ii) enhanced MMPBSA calculations based on the Temperature Replica Exchange method and (iii) prediction based on Rosetta force field – on the same dataset of point (ii) – and (iv) calculation of the PMF using an umbrella sampling algorithm. The key result is that all methods based on MD simulations can be used to produce good qualitative estimate of the binding energy differences between different antibodies, even if the quality of prediction depends on the methodology adopted. Furthermore, energy determinations calculated with MMPBSA are more compatible with the experiments than those obtained with Rosetta. However, an important shortcoming of MMPBSA is the large variance of the results, even within a well-equilibrated and homogeneous simulation. For this reason, it is essential to obtain an accurate sampling of the configuration space.

The reliability of the different methods also appears to be system specific. Indeed, PMF calculations based on umbrella sampling have shown high accuracy in other systems [23], but underperformed in our case. We can hypothesize that the specific characteristics of the system contribute to the unexpectedly low quality of predictions obtained from the Rosetta force field or from the PMF determination. Since the peptide interacts with the antibody via two hydrophobic residues, the two molecules separate abruptly under the external force, making the sampling of intermediate positions more difficult.

Another point we analyzed was the effect of the sampling size vs the quality of predictions. Both RE-MMPBSA and PMF required long simulations for the evaluation of each binding affinity (6.4 μs and 300 ns, respectively). Still, the results correlate more poorly (R^2^ = 0.31 and R^2^ = 0.19, respectively) than those presented in our previous work, which required 50 ns (10 replicas of 5 ns simulations) for each calculation (R^2^ = 0.57). Consistently with our previous observation, shorter simulations produced better results, and correlation significantly improved when we eliminated the second part of the trajectories (R^2^ = 0.57).

This counterintuitive finding - that longer simulations do not always translate into more accurate predictions - likely stems from several factors. Atomistic force fields could show some limitations in reproducing protein-protein interactions correctly [33]. Furthermore, longer simulations may drift from the true conformational ensemble or become trapped in local minima that are nearly optimal but not representative of the native binding state. This observed effect complicates the development of standardized protocols, as optimal simulation length appears to require empirical determination for each system type.

From a practical standpoint, however, this has very important implications, because we have proven that short simulations are sufficient to reproduce experimental results qualitatively, and therefore can be used effectively in a screening process. Based on our results, the best strategy for rapid screening of many different possible binders to a given target is the one originally proposed in [20]. It consisted of running 10 different simulations of 5 ns equilibrium dynamics. From the second half of these simulations, we extracted configurations every 100 ps and calculated the binding affinity as the average of the binding free energies obtained with MMPBSA. This method outperforms methodologies based on effective potentials (such as PRODIGY [18] or Rosetta [22]), while remaining fast enough for rapid screening.

We also observed some cases in which different simulation methods tend to agree on the qualitative ranking of the antibodies but disagree with experimental evidence. Even if we have treated experiments as the ground truth so far, we must recognize that experiments come with errors as well, and good leads may be lost in the process because of experimental uncertainties.

Given these advantages, computational methods are emerging as a valid complement to, and for screening purposes, a powerful alternative to experimental approaches. Their financial cost is significantly lower, since a high-performance workstation is far less expensive than the laboratory infrastructure required for experimental binding assays, and no consumables are needed. The time cost may also be favorable: with modern GPUs able to simulate ∼300 ns/day for systems of this size, the proposed protocol allows testing 3–5 antibodies per GPU per day. In contrast, experiments often require several time-consuming steps, such as antibody expression and purification, so that collecting equivalent data may take days to weeks.

Another important advantage of computational methodologies is their potential to optimize antibody binders against particularly challenging targets, such as connexins or ASIC channels [34,35]. These proteins have been proven difficult to characterize with standard experimental assays but can be explored effectively with the computational pipeline described here.

## 4. Conclusions

Our results prove that binding affinities predictions based on MD simulations – when paired with an extensive sampling of the configuration space – provide a valuable tool for screening of antibodies, and this can certainly be extended to small molecules and peptides alike. The key to success is the sampling of the configuration space, which allows us to account for the entropic contribution to the binding free energy. However, since longer simulations can actually reduce predictive power, careful empirical calibration of simulation length is required. While the computational cost of extensive sampling is relatively high, the growing availability of computational power is making this approach increasingly feasible, and it may become the gold standard in drug design pipelines. For immediate practical applications in screening, however, the simpler protocol of running multiple short simulation replicas provides an effective compromise between accuracy and computational feasibility.

## 5. Materials and Methods

### 5.1 Antibodies

The antibodies considered in this paper have been characterized experimentally in previous work [20,26]. To simplify the notation in this work, the original antibody abN48 has been renamed AbWT, and its variant has been renamed Ab02, …, Ab38 from abN48-2, …, abN48-38.

### 5.2 General information about MD simulations

Simulations were conducted using the Gromacs 2023.2 package [36], using the Amber14SB force field [37] and the TIP3p water model, following protocols similar to our previous works [38,39] and explained hereafter briefly.

### 5.3 Initial equilibration

Molecular systems composed of antibodies and antigens were simulated under periodic boundary conditions. Each system was solvated using TIP3P water, with the addition of 0.15 M KCl to mimic physiological conditions. After neutralization, energy minimization was performed to prevent steric clashes and resolve potential energy conflicts within the system. Equilibration followed, involving two short MD Simulations: first in the NVT volume ensemble, then in the NPT ensemble to equilibrate side chains. Each simulation lasted 100 ps, with positional restraints applied to the alpha carbon atoms, and the LINCS algorithm was adopted to constrain the H-bonds and allow an integration timestep of 2 fs. constraints applied to hydrogen bonds to ensure structural integrity.

### 5.4 Restricted temperature replica exchange molecular dynamics

To reduce the computational cost of the calculations, we limited the T-REMD strategy to the interface between the antibodies and peptides. Antibody atoms outside the interacting shell, defined by all the residues within 15 angstroms with respect to the interacting peptide, were harmonically restrained at the starting position with a force constant of 10 kJ mol^-1^ in the three spatial dimensions during the T-REMD simulations. Specifically, 64 parallel simulations were run in NPT standard conditions for 100 ns, restraining the atoms outside the interacting shell. The temperature distribution ranged from 300 to 380 K, and an exchange trial between adjacent replicas was performed every 1,000 steps of 2 fs. The solvation of the systems in an octahedral box was done using the TIP3P water model with a 1.1-nm distance to the border of the molecule. A concentration of 150-mM KCl and additional ions was used to neutralize. To control temperature and pressure, the Berendsen algorithm was applied, and electrostatic interactions were treated using the Particle Mesh Ewald method [40]. Water molecules were first relaxed by energy minimization and 10 ps of simulations at 300 K, restraining all the protein and peptide atomic positions with a harmonic potential. Then, the systems were heated up gradually to 300 K in a six-step phase starting from 50 K, and the final run was performed restraining the atomic position of the residues out of the binding region

### 5.5 Molecular Mechanics Poisson-Boltzmann Surface Area (MMPBSA)

Binding free energy (Δ*G*) were computed using the MMPBSA method within the single-trajectory approximation framework [15]. MMPBSA is a computational approach that estimates binding affinities by combining molecular mechanics (MM) energies with continuum solvation models. The method decomposes the free energy into gas-phase molecular mechanics terms computed from the forcefield parameters (bonded, van der Waals, and electrostatic interactions), polar solvation energy calculated using the Poisson-Boltzmann equation, and non-polar solvation energy estimated from solvent-accessible surface area (SASA). The final estimates for the binding free energies were calculated as averages over configurations extracted from the T-REMD trajectories every 100 ps in the equilibrated portions of the simulations (20-50 ns or 20-100 ns). The binding free energy was calculated as Δ*G*= *G*_*complex*_ − (*G*_*peptide*_ + *G*_*antibody*_) where *G*_*complex*_ represents the free energy of the protein-ligand assembly, and *G*_*peptide*_ and *G*_*antibody*_ correspond to the free energies of the unbound peptide and antibody, respectively. Calculations were carried out using the gmx_MMPBSA tool [41,42].

### 5.6 Calculation of binding energies using Rosetta force field

The input PDB file was first processed using Rosetta’s *score_jd2* application to repair the structure and ensure compatibility with Rosetta. Following this, Rosetta’s *InterfaceAnalyzer* was employed with key parameters enabled to analyze protein-protein interactions. The *-pack_input true* option was used to optimize side-chain conformations at the interface, while *-pack_separated true* facilitated the calculation of binding energy (*dG_separated*) by repacking separated chains to model unbound states. The standard *ref2015* energy function [22] was applied for scoring. Additionally, interface packing quality was assessed by enabling the *-compute_packstat true* parameter to evaluate packing statistics. This approach ensured a comprehensive analysis of the protein interface while maintaining structural integrity.

### 5.7 Steered molecular dynamics

Initial configurations for the umbrella sampling simulations were obtained using a non-equilibrium pulling simulation (steered MD Simulation). A harmonic potential (2000 kJ/mol·nm2) was applied along the z-axis of the system between the centers of the two pull groups: pull group 1 on the antibody and pull group 2 on the peptide. Pull group 1 consisted of 13 atoms selected from alpha carbon atoms on the backbone, spanning positions 60 to 100 along the z-axis, while pull group 2 contained 7 alpha carbon backbone atoms. Constraints on hydrogen bonds were applied to maintain structural integrity during the simulation. The MD simulation was run for 3 nanoseconds with a pulling rate of 2 nm/ns. The Particle-Mesh Ewald method was used for handling long-range electrostatics [40].

### 5.8 Umbrella sampling

The PMF was reconstructed by applying the umbrella sampling method [16,24,25]. The reaction coordinate (ζ) corresponded to the distance between the center of mass of the two pull groups defined above. The distance between the various umbrella sampling windows was set to 0.3 Å, totaling about 100 windows to cover the whole reaction coordinate. Each window was simulated for 2ns. The umbrella biasing potential was set by a quadratic potential with K = (2000 kJ/mol·nm^2^). Finally, the Weight Histogram Analysis Method [43] was used to obtain the PMF for each simulation.

## Statistical analysis of autocorrelation

As configurations are extracted from molecular dynamics trajectories, the samples are autocorrelated, leading to an overestimation of the effective sample size and an underestimation of the true statistical uncertainty. To address this issue, we used the PyMBAR Python library to analyze the statistical properties of each umbrella sampling window and estimate the number of effectively uncorrelated samples, which have been used to calculate averages and standard deviations.

## Aknowledgment

We thank Dr. Luigi Vitagliano for helpful discussion.

## Founding sources

This work was partially supported by Xi’an Jiaotong-Liverpool University: Research Development Fund RDF-23-01-026 to FZ and RDF-24-01-073 to KCC, by Zhejiang University: Fundamental Research Funds for the Central University (226-2024-00123) to DB, and by the National Key R&D Program of China (2024YFA1306400, 2021YFA1201200, 2024YFA1307500), the National Center of Technology Innovation for Biopharmaceuticals (NCTIB2022HS02010), Shanghai Artificial Intelligence Lab (P22KN00272), the Starry Night Science Fund of Zhejiang University Shanghai Institute for Advanced Study (SN-ZJU-SIAS-003) to RZ.

Simulations were carried on at the Xi’an Jiaotong-Liverpool University HPC, the Institute of Quantitative Biology of Zhejiang University HPC and the CINECA Supercomputing center (ISCRA-C projects: C: MDpapt - HP10CL0DAJ, MD-dir2 - HP10CZVX8N, RNAmsd - HP10CGPJAR, dad - HP10CU5FCP, RnaMod - HP10C559YV).

## Declaration of generative AI and AI-assisted technologies in the writing process

During the preparation of this work the authors used ClaudeAI, ChatGPT and Grammarly in order to improve readability and correct typos or grammar mistakes. No part of the content has been generated by AI tools. After using these tools, the authors reviewed and edited the content as needed and take full responsibility for the content of the published article.

## Figures

**Supplementary Figure 1.**
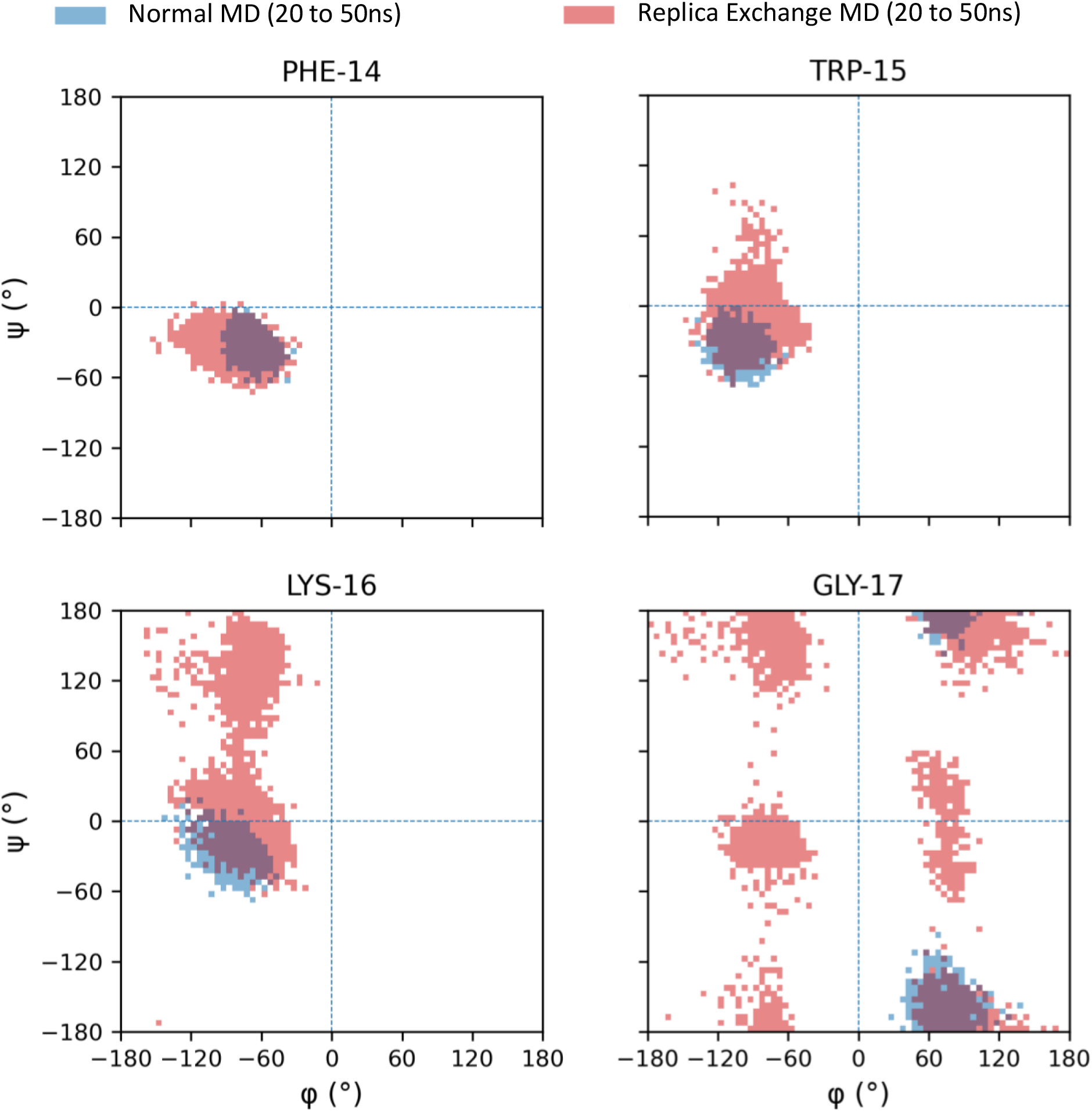
Ramachandran plots for normal MD simulations vs T-REMD simulations. The four graphs report the values of the Ramachandran angles for four critical peptide residues (PHE14, TRP15, LYS16 and GLY17). Blue points represent the values for standard MD simulations, while red points for T-REMD. It is evident how in the second case the peptide can explore a vaster configuration space.

**Supplementary Figure 2.**
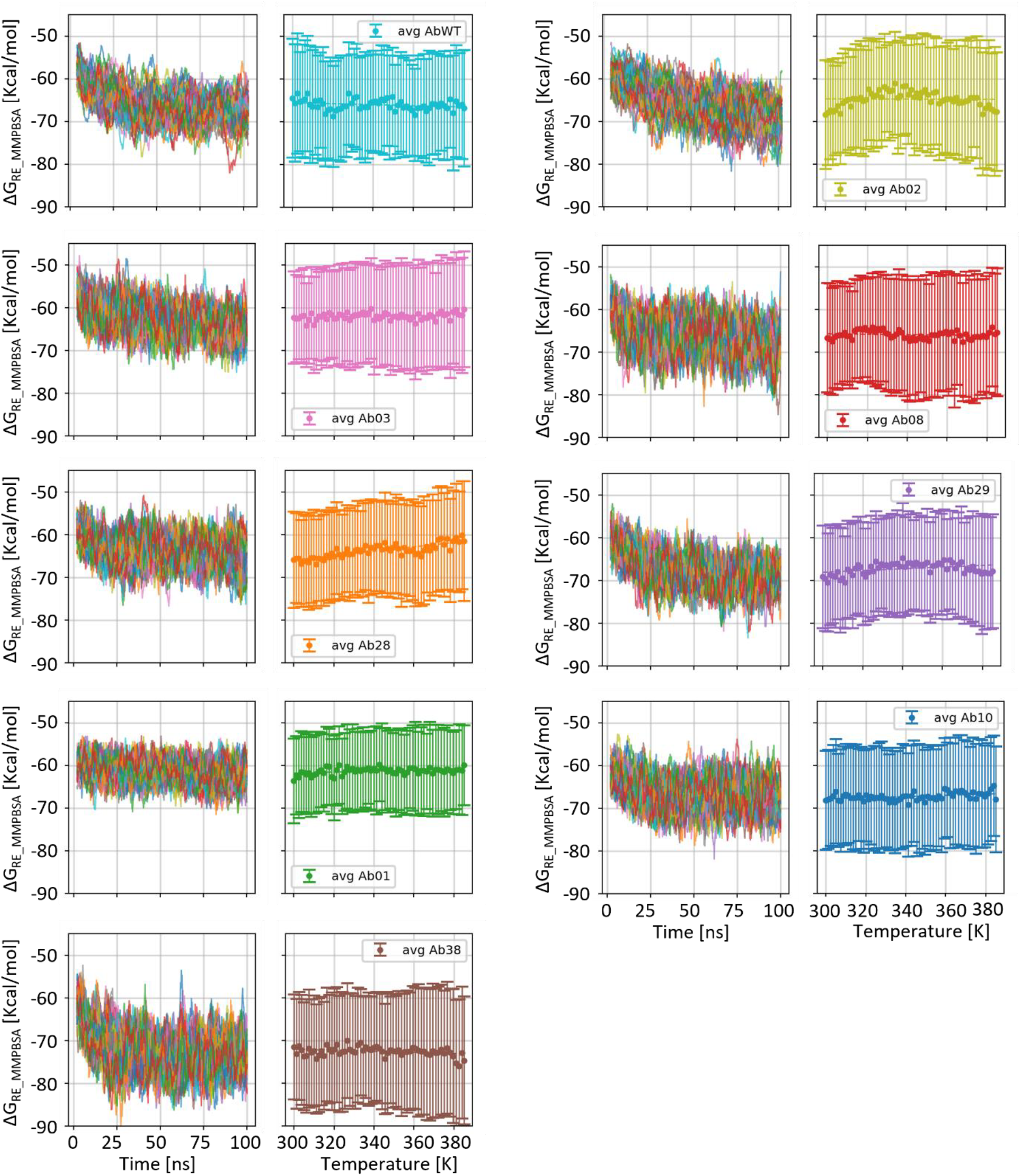
Temperature dependence of *ΔG* in the T-REMD simulations. For each antibody, we report the *ΔG*values as a function of time for all the different replicas (panels on the left of each subfigure) and the average and standard deviation of the binding affinities calculated in the time period 20-50 ns for all the temperatures considered (panels on the right of each subfigure). Different antibodies thermalize at different times, but generally a plateau is reached after 20 ns. It can also be noticed that temperature dependence is very weak or absent in most cases.

